# A preliminary ecological profile of Kyasanur Forest disease virus hosts among the mammalian wildlife of the Western Ghats, India

**DOI:** 10.1101/2020.01.30.927939

**Authors:** Michael G. Walsh, Siobhan M. Mor, Hindol Maity, Shah Hossain

**Affiliations:** The University of Sydney, Faculty of Medicine and Health, Marie Bashir Institute for Infectious Diseases and Biosecurity, Westmead, New South Wales, Australia; The University of Sydney, Faculty of Medicine and Health, Westmead Institute for Medical Research, Westmead, New South Wales, Australia; Prasanna School of Public Health, Manipal Academy of Higher Education, Manipal, Karnataka, India; University of Liverpool, Faculty of Health and Life Sciences, Institute of Infection and Global Health Liverpool, Merseyside, United Kingdom; The University of Sydney, Faculty of Science, School of Veterinary Science, Camperdown, New South Wales, Australia; Manipal Academy of Higher Education, Manipal, Karnataka, India

**Keywords:** Kyasanur Forest disease, tick-borne disease, infection ecology, epidemiology, reservoir host

## Abstract

Kyasanur Forest disease (KFD) is one of India’s severe arboviruses capable of causing prolonged debilitating disease. It has been expanding beyond its historical endemic locus at an alarming rate over the last two decades. The natural nidus of this zoonosis is located in the monsoon rainforest of the Western Ghats, India, which is one of the world’s most important biodiversity hotspots. Definitive reservoir hosts for KFD virus (KFDV) have yet to be delineated, and thus much of the infection ecology of this virus, and its consequent transmission dynamics, remains uncertain. Given its unique biogeographical context, identifying ecological parameters of KFDV relevant to the virus’ epidemiology has been complex and challenging. The challenge has been exacerbated by diminished research efforts in wildlife surveillance over the last two decades, coinciding with the expansion of the range of KFD across the region. The current investigation sought to define a preliminary ecological profile of KFDV hosts based on their life history and feeding traits to aid in re-establishing targeted wildlife surveillance and to discern those ecological traits of wildlife hosts that may improve our understanding of KFD epidemiology. The importance of fast-living among KFDV hosts was of special interest with respect to the latter aim. We compared mammalian traits between host and non-host species using general additive models and phylogenetic generalised linear models. This study found that both body mass and forest forage were strongly associated with mammalian host infection status, but that reproductive life history traits were not. These findings will help in structuring ecologically based wildlife surveillance and field investigations, while also helping to parameterise novel epidemiological models of zoonotic infection risk that incorporate species functional traits in a region where biogeography, landscape ecology, and community ecology manifest extraordinary complexity, particularly under growing anthropogenic pressure.

## Introduction

Kyasanur Forest disease (KFD) is a severe tick-borne infection endemic to the Western Ghats of South India. In humans, clinical symptoms are biphasic, with the initial acute phase typically manifesting as flu-like symptoms as well as severe muscle pain, vomiting, diarrhea and bleeding, followed by an afebrile period during which most patients convalesce. A second phase presenting with meningoencephalitis affects approximately 10-20% of patients one to two weeks following the resolution of the acute phase (Pattnaik, 2006; Shah et al., 2018). The disease is caused by the flavivirus, Kysanaur Forest disease virus (KFDV), which is transmitted primarily by the forest tick, *Haemaphysalis spinigera*. Incidence has increased annually over the last two decades (Mourya and Yadav, 2016; Pattnaik, 2006), and has expanded rapidly beyond its original foci of the Kyasanur Forest area of Karnataka State in recent years (Figure 1) (Mourya and Yadav, 2016; Pattnaik, 2006; Shah et al., 2018). These observations are supported by tick field surveys which demonstrate increasing infection risk (Naren babu et al., 2019). Recent work has shown that the landscape epidemiology of KFD is strongly influenced by the loss of forest habitat in the Western Ghats, and that areas with high mammalian species richness are particularly suitable (Walsh et al., 2019). Combined, these findings suggest that anthropogenic pressure on wildlife habitat may be inadvertently increasing human exposure to KFD reservoirs and their tick vectors.

**Figure 1.**
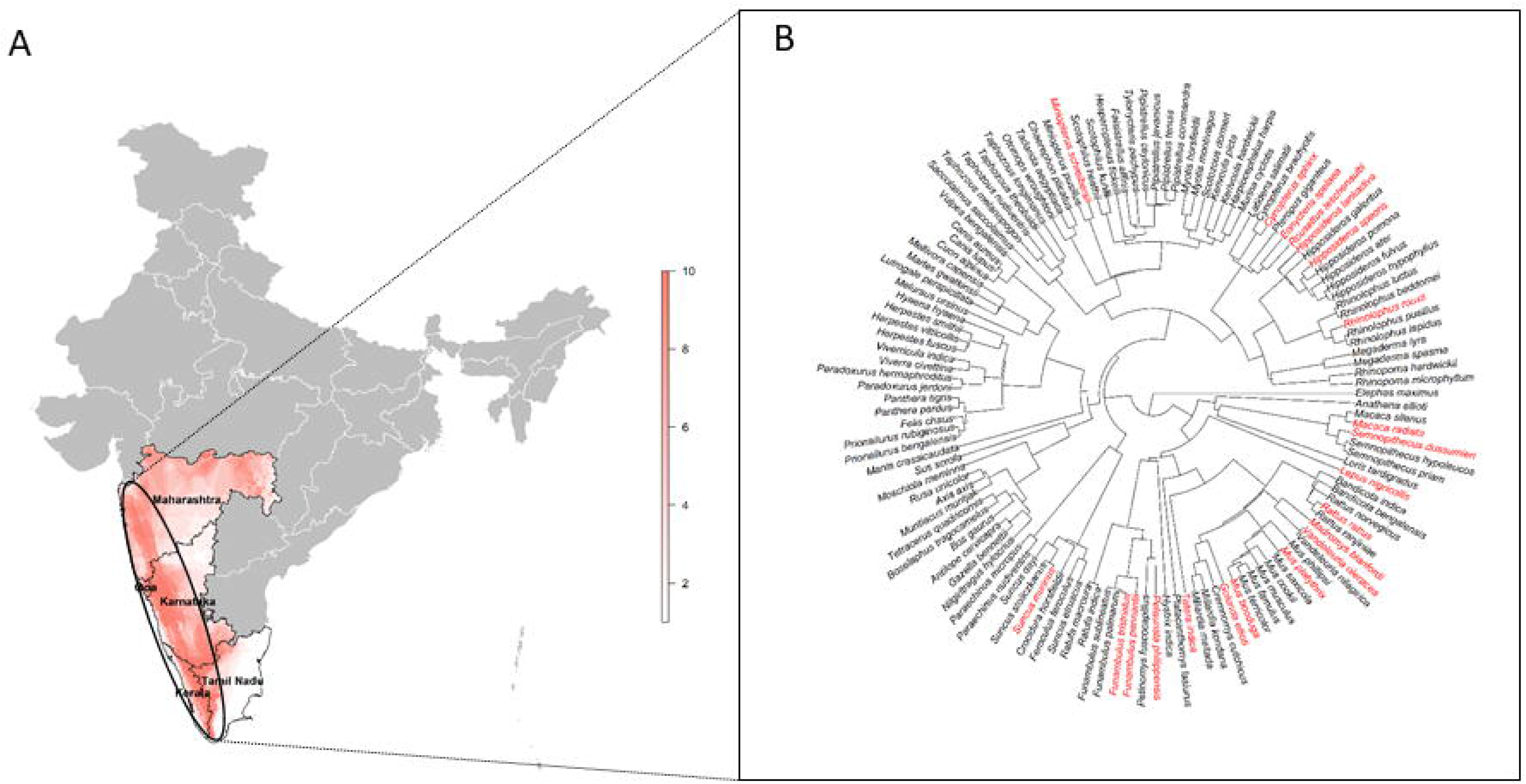
The Western Ghats states of South India highlighted with A) deciles of mammalian biodiversity across each state, generally, and within the Western Ghats region (oval), specifically, and B) a phylogenetic tree of mammalian species present in the Western Ghats, with documented Kyasanur Forest disease virus hosts identified in red. This map is used only for the purposes of representing mammalian species and does not reflect the authors’ assertion of territory or borders of any sovereign country including India. All maps created in R (v. 3.3.1).

Nevertheless, we currently have a limited understanding of the ecology of and transmission dynamics in the virus’ wildlife hosts. In the years after KFDV was first identified in 1957 there was a brief surge in research that attempted to identify the virus’ wildlife reservoir in the Kyasanur Forest and adjacent areas of the larger Bandipur Forest range (Pattnaik, 2006). Several mammalian species were identified as possible reservoirs, primarily among bats and rodents (Pattnaik, 2006). However, the research slowed considerably by the 1970s and ultimately stopped by the early 1980s, before definitive reservoirs were established as those species (or communities of species) capable of maintaining virus circulation indefinitely, or as those species capable of providing a source of virus for tick-borne human infection irrespective of the species’ capacity to maintain virus circulation (Haydon et al., 2002)‥ As such, we remain largely ignorant of enzootic and epizootic cycling of KFDV in its natural nidi and the subsequent effects of such wildlife circulation on zoonotic transmission.

Recent work on other pathogens has shown that life history may modulate infection systems that have high potential for spillover (Huang et al., 2013; Ostfeld et al., 2014). In particular, “fast-living” species are associated with reservoir competence (Brunner et al., 2008; Ostfeld et al., 2014), a finding that is hypothesised to be immunologically-mediated (Johnson et al., 2012). Fast-living species are characterised by a shorter lifespan and higher reproductive output coupled with lower early development investment. Examples of a fast-living reproductive profile include more litters, shorter inter-birth intervals, shorter length of gestation, and earlier weaning compared to slow-living species (Molles, 2015). Strong evidence in support of the relationship between fast-living and host competence, which we define here as a host’s potential to generate new infections in other hosts (Downs et al., 2019), was provided by a comprehensive survey of all mammalian host–pathogen systems (Plourde et al., 2017). However, the reverse has been demonstrated for some zoonotic arboviruses (Walsh and Mor, 2018; Walsh, 2019) and Ebolavirus (Schmidt et al., 2019), which demonstrated slow-living to be associated with mammalian hosts. Trait-based approaches have also been used effectively for reservoir species prediction both in specific zoonosis systems (Han et al., 2016; Plowright et al., 2019) and for general zoonoses within specific taxa (Han et al., 2015).

The current investigation sought to identify all mammalian species previously recorded as susceptible to infection with KFDV and to compare these to all other species present in the Western Ghats with respect to ecological and life history traits. In this way we sought to delineate a trait-based profile for wildlife hosts that can serve to 1) inform KFDV surveillance among wildlife using a new ecologically-targeted approach, and 2) identify patterns of species-environment interaction that may contribute to the epidemiology of KFDV.

## Materials and Methods

### Data sources

One hundred and thirty-five mammalian species were previously documented in the Western Ghats region (Nameer et al., 2001). However, this was recently updated by the Zoological Survey of India, which now reports 133 mammal species present representing the Chiroptera, Rodentia, Carnivora, Artiodactyla, Eulipotyphla, Primates, Lagomorpha, Scandentia, Pholidota and Proboscidea(Venkataraman et al., 2018). Of these, 21 have been documented as susceptible to KFDV infection by serology or virus isolation (Bhat et al., 1979, 1978; Boshell et al., 1968; Goverdhan and Anderson, 1981, 1972; Pavri and Singh, 1968, 1965; Rajagopalan et al., 1969a, 1969b; Sreenivasan and Bhat, 1976, 1977; Sreenivasan et al., 1979; Webb, 1965; Work, 1958). The remaining species with no documented infection were classified as undetermined KFDV status. The Western Ghats region, its biodiversity, and a phylogenetic tree of its mammalian species are presented in Figure 1. The phylogenetic tree was obtained from the VertLife project (Upham et al., 2019). A table of all listed species along with their host susceptibility status is provided in Figshare. It must be stressed that screening by the multiple modalities described precludes qualitative designation of species as reservoir, maintenance, or amplification hosts. The hosts can only be designated as susceptible to infection and we concede the limitations of the generic inference derived. Among those species identified as susceptible to KFDV infection, 10 were rodents, 7 were bats, 2 primates, 1 hare (Lagomorpha), and 1 shrew (Eulipotyphla) (Figure 1). In order to correct for potential bias introduced by differences in reporting effort across species, the number of published studies for each species in the Web of Science (WoS) database was used to quantify reporting effort. This was then included as an additional covariate in the model (see statistical analysis below), as done in previous studies (Nunn et al., 2003; Olival et al., 2017). For species whose range included the Western Ghats but was not endemic to the region, only those studies conducted in the region were included from the WoS. Ideally it would be preferable to evaluate these models in only those species that have been surveyed for KFDV. However, this was not feasible due to the previously described absence of wildlife KFDV surveillance over the last several decades. Nevertheless, as an additional sensitivity analysis the data were restricted to only those species (n = 54) that have been previously surveyed for viral infection as a proxy for species’ studiedness in relation to viral surveillance in wildlife. This reduced dataset was based on the mammal-virus association database compiled by Olival and colleagues (Olival et al., 2017). Model outputs thus attempted rigorous control of reporting effort for what has otherwise been a poorly surveyed zoonotic virus in wildlife hosts.

Two databases were used to quantify the ecological and life history traits of the Western Ghats mammal species. The Elton Traits database was used to obtain species’ diet composition, foraging strategy, and circadian patterns of activity (Wilman et al., 2014). In this dataset diet composition comprised several individual plant food sources (fruit, nectar, seeds, and grasses, leaves, and other ground-based plants), which were aggregated here to characterise the overall proportion of the diet derived from plants. Similarly, the proportion of the diet derived from vertebrates was aggregated across prey endothermic vertebrates (mammals and birds), ectothermic vertebrates (reptiles and amphibians), fish, and scavenged vertebrates, i.e. the scavenged carcasses of any of the above vertebrates killed by other animals. The third dietary category considered here was the proportion of the diet derived from invertebrates. Foraging strategy comprised four categories: ground-based, scansorial (climbing foraging), arboreal (foraging in trees), and aerial. These four categories were constructed as a numeric ordinal variable from 1 to 4, with increasing values corresponding to increasing foraging off the ground. This construct was based on the hypothesis that mammals that spend more time foraging on the ground would be more accessible to tick vectors. The PanTHERIA database was used to obtain life history traits and body mass (Jones et al., 2009). Variables with low missing data (≥ 70% complete) and which capture the spectrum of life history from fast- to slow-living (Bielby et al., 2007) were included in the analysis, namely maximum longevity, sexual maturity age, gestation length, inter-birth interval, weaning age, and litter size. Adult body mass and neonatal body mass also demonstrated low missingness among these species in the PanTHERIA database and were used to construct the new life history metric of mass gain, adjusted for adult body mass: (adult mass – neonate mass)/adult mass. Any remaining missing trait data for these selected traits were imputed using a random forest machine learning algorithm, which has previously been shown to be robust imputation for these data (Plourde et al., 2017; Stekhoven and Buhlmann, 2012). The rfImpute function in the randomForest package was used to implement the algorithm (Liaw and Wiener, 2002). The traits are presented by taxonomy in S1 Figure 1.

### Statistical Analysis

Blomberg’s K was used to test for the presence of a phylogenetic signal for any of the ecological or life history traits (Blomberg et al., 2003). All K were well below the conservative 0.5 threshold (S2 Table 1), which was strong evidence of the absence of phylogenetic correlation among these traits. As such, the use of a model framework accounting for phylogenetic correlation structure, such as a phylogenetic generalised linear model (PGLM) was deemed unnecessary. We therefore used generalised additive models (GAMs) to evaluate non-linear relationships between KFDV infection status and species’ traits. These models fit multiple basis functions for each smoothed covariate, thus allowing the covariate to vary in a non-linear fashion across its distribution with respect to an outcome (Wood, 2017, 2004). Because each covariate is represented by multiple parameter estimates rather than a single summary parameter estimate (i.e. as with a regression coefficient in a GLM), the relationships between covariates and outcomes must be graphed to show how the outcome varies non-linearly over the distribution of the covariate. Categorical covariates, however, are fit as non-smoothed fixed effects and thus retain a single parameter estimate summarising their relationship with the outcome. Since the outcome modelled here is dichotomous (KFDV infection status), the binomial family was used as the link function in these GAMs. A principal component analysis was used to construct orthogonal life history metrics from the PanTHERIA data due to the high correlation between the life history traits among the mammals of the Western Ghats. The principal components were then included as the life history trait covariates in the multiple GAMs along with the Elton traits. The full model comprised all selected Elton species traits (forage strategy and proportion of the diet derived from plants), life history trait principal components (PC), body mass, and reporting effort (correlations among these variables was low; all *r* < 0.6). The best GAM was determined using double penalty smoothing selection (Marra and Wood, 2011).

Three sensitivity analyses were performed to test model validity. First, to account for any residual influence of phylogenetic correlation operating within the relationships quantified by the GAM, the GAM was compared to a PGLM to fully evaluate the influence of the phylogenetic signal across the two model structures. Second, an alternate GAM was fit with mass-corrected life history to explore whether the relationships between life history traits and infection status might scale with body mass. Third, as described above, the GAM based on the complete dataset of mammalian species of the Western Ghats was verified with a reduced dataset comprised of only those mammals with documented viral surveillance. The R statistical platform v. 3.6.1 was used for all analyses (R Core Team, 2016). Phylogenetic correlation was quantified using the ape package (Paradis et al., 2004; Popescu et al., 2012), while Blomberg’s K was estimated using the phylosig function in the phytools package (Revell, 2012). All GAMs were fit using the gam function in the mgcv package (Wood, 2017). The PGLM was estimated using the phyloglm function in the phylolm package(Tung Ho and Ané, 2014).

## Results

Bivariate comparisons of each trait by host status are presented in Figure 2 and S3 Table 2. Diet source and adult body mass showed the strongest associations with KFDV infection status, wherein hosts were smaller and obtained the majority of their diet from plants. The crude comparison of individual life history traits showed a tendency toward fast-living, with hosts demonstrating shorter lifespans and inter-birth intervals, and younger weaning and sexual maturity (Figure 2). The principal components analysis identified three factors that cumulatively explained 90% of the variance among these highly correlated life history traits and were thus included in the GAMs to represent the fast- to slow-living spectrum (S4 Table 3A). The first of the three factors was positively correlated with slow living across all life history traits, as was the second with the exception of a negative correlation with slow living in gestation length. The third factor, conversely, was negatively correlated with slow living across these traits, with the exception of weaning age (S4 Table 3B).

**Figure 2.**
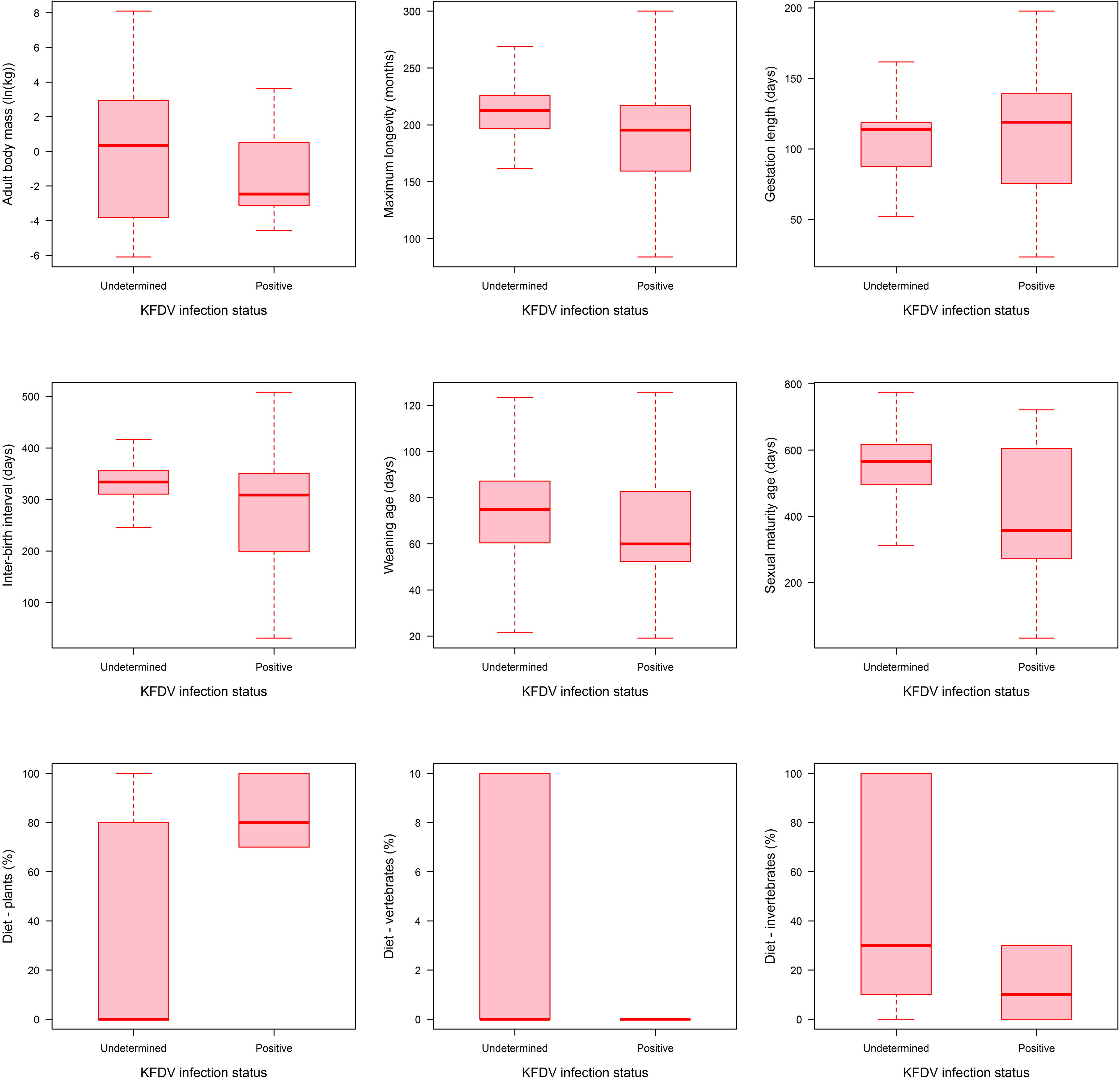
Comparison of ecological and life history traits among the mammalian species of the Western Ghats, stratified by documented Kyasanur Forest disease virus host status.

The best fitting GAM selected by double penalty smoothing (Model 1) is presented in Table 1. This GAM comprised only two traits, body mass and a predominantly plant-based diet, which demonstrated strong non-linear relationships with the probability of being a KFDV host. Life history and foraging strategy were no longer associated with infection status after accounting for body mass and diet, while reporting effort showed no association with infection status bivariately (S3 Table 2) or in the multiple GAM. Comparing the residual deviance (89.0) from the model to the null deviance (115.0) further showed the GAM to be a good fit to the data (p>0.99). In addition, the residuals from the GAM were further tested using Blomberg’s K and identified no phylogenetic signal (K = 0.04). An increasing proportion of the diet derived from plants was associated increasing probability of being a host (Figure 3). Body mass indicated a very high probability of being a host among smaller species, but then dropped precipitously among the larger species. To account for any residual influence of phylogenetic correlation the first sensitivity analysis replaced the GAM with a PGLM (Model 2 in Table 1). This model identified the same relationships with body mass and plant-based diet, although the associations were somewhat attenuated. Given the strong association with body mass, it was of further interest to explore whether the relationships between life history traits and infection status might scale with body mass. The alternate GAM from this second sensitivity analysis (S5 Table 4) confirmed the lack of association between life history and infection status even when the former was scaled to body mass. The third sensitivity analysis based on the reduced dataset of virus-surveyed species produced the same best fit model, comprising body mass and plant-based diet. Moreover, the functional relationship was very similar albeit with wider confidence limits as expected due to the reduced sample size (S6 Figure 2).

**Table 1.** Generalised additive model (GAM) and phylogenetic generalised linear model (PGLM) of Kyasanur Forest disease virus infection status and mammalian species’ traits.

| Model | Species trait | p-value* |
| --- | --- | --- |
| Model 1 – GAM | Adult body mass (ln(kg)) | 0.0005 |
|  | Diet – plant (%) | 0.00002 |
| Model 2 – PGLM | Adult body mass (ln(kg)) | 0.02 |
|  | Diet – plant (%) | 0.001 |

\*p-values refer to the association between each species' trait and infection status, not the overall fit of the model.

**Figure 3.**
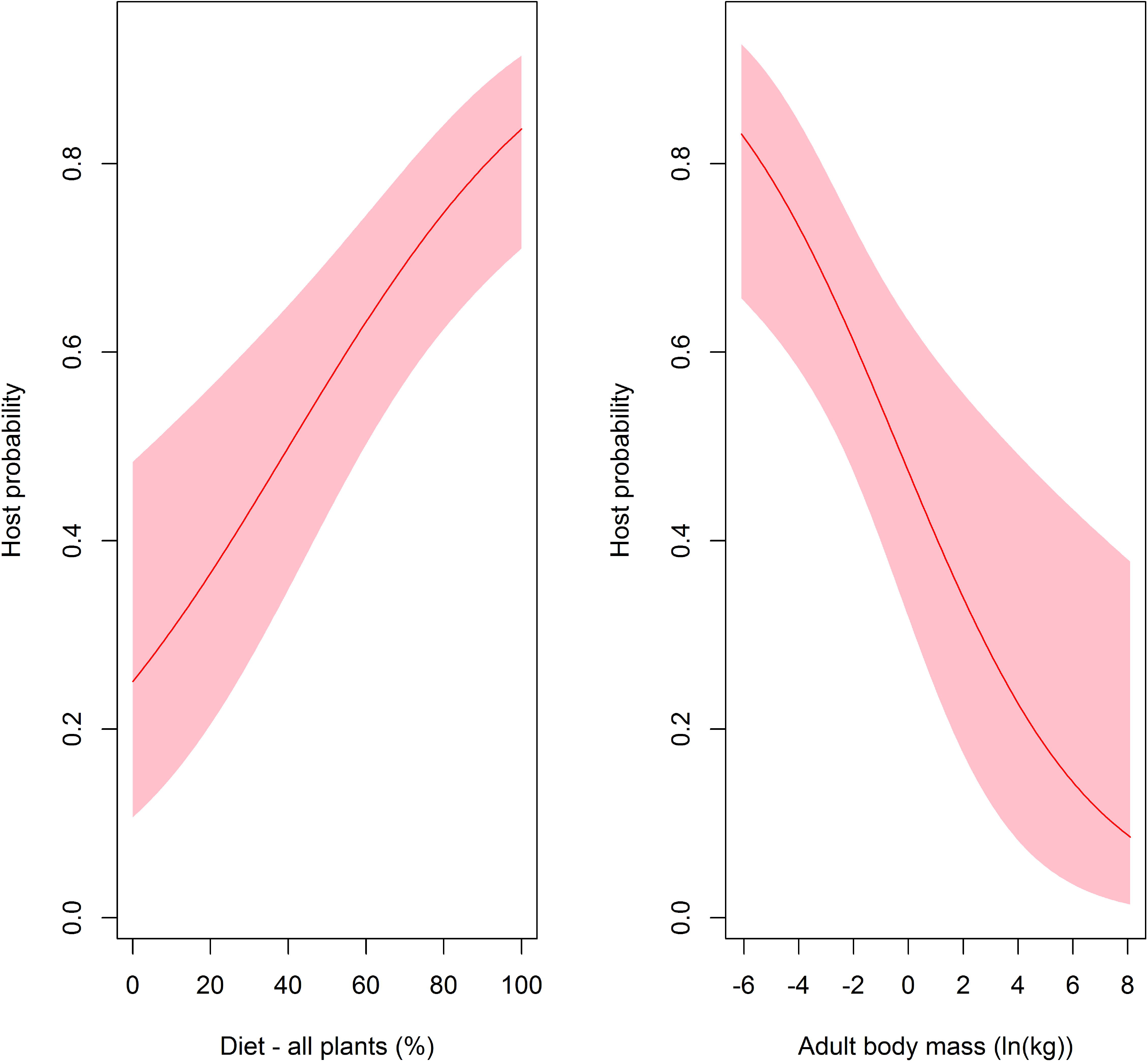
The nonlinear relationships between Kyasanur Forest disease virus host probability and plant-based diet and body mass as derived from the best fitting generalised additive model. Shaded areas represent the 95% confidence limits of the nonlinear function. Diet here represents the percentage of the species’ diet derived from plants and body bass is on the natural log scale.

## Discussion

This study presents a preliminary ecological profile of mammalian hosts of KFDV. The findings showed that host species tended to be small and to derive more of their dietary intake from plants. As such, this preliminary ecological profile of KFDV hosts highlights species’ use of forest forage, which may be particularly relevant for the implications of habitat conservation on public health, and size, which may reflect a common pattern of hypometric scaling among reservoir hosts of tick-borne pathogens.

Kyasanur Forest disease virus hosts derived a substantively greater proportion of their diet from plants compared to non-hosts. Moreover, the gradient of host probability increased as the plant-based diet increased. This relationship suggests an important nutritional dependence of KFDV hosts directly on forest plant resources, i.e. primary production. When such resources are lost due to deforestation (Jha et al., 2000), the subsequent displacement of animals may lead to foraging in novel, anthropogenic ecotones located across the transition from forest to human habitation or agriculture, potentially exposing humans to their ticks and arboviruses (Ajesh et al., 2017; Naren babu et al., 2019; Sadanandane et al., 2018). This may be reflected in the shifting of the nidus of KFD from Sagar and Soraba Taluks in Shimoga district in Karnataka northwards into Goa and Maharashtra and southwards into Thirthahalli Taluk in Karnataka, Wayanad district in Kerala, and Nilgiri in Tamil Nadu, all of which comprise the Bandipur Forest range (Walsh et al., 2019). Novel interspecific interactions and increasing wildlife-human conflict have previously been documented specifically in forest fringe areas of the Western Ghats (MADHUSUDAN, 2004), such that the potential for zoonotic transmission is already well established. Moreover, the importance of human pressure on natural landscapes has been shown in similar contexts of emerging zoonoses of wildlife origin (Allen et al., 2017).

Body mass manifested significant influence on mammalian host status. Several mechanisms by which body mass can modulate infection transmission dynamics have been articulated with respect to the scaling of host competence, in particular (Downs et al., 2019). Scaling, in this context, refers to the degree to which a particular trait (e.g. KFDV competence) is geometrically similar across the spectrum of body size. With isometric scaling, the trait of interest does not differ between large and small animals, whereas with allometric scaling the trait differs by body size. Competence can be thought of as a host’s potential to generate new infections and operates as a function of the species’ exposure, susceptibility suitability and transmissibility of the pathogen (Downs et al., 2019). Downs et al. showed distinctive patterns of allometric scaling with body size, and the direction (hypermetric vs. hypometric scaling) can differ for different pathogens. A distinct hypometric scaling has been shown for other tick-borne pathogens wherein smaller mammals demonstrated a greater likelihood of being competent hosts (Barbour et al., 2015; Ostfeld et al., 2014). It is important to note, however, that the current study was not able to assess host competence directly and so the current findings can say nothing definitive about the allometric scaling of competence with respect to KFDV. Rather, the findings suggest that this relationship could provide a useful framework for future field investigations. Species population density is also important to consider with respect to species body mass, given that allometric scaling of host traits may operate through the product of individual size and population density. For example, even systems demonstrating hypermetric scaling have shown that transmission potential for species of smaller mass can be greater because these species also manifested higher population density, which translated to higher transmission potential (Downs et al., 2019). The current study was not able to adequately assess the contribution of population density to the KFDV infection status due to the significant missing data for this variable in PanTHERIA database, thus precluding it from the data imputation. Further work is required to evaluate the synergy between species individual mass and species population density.

Fast-living was hypothesised to be more prominent among KFDV host species. Faster-living mammal species were more likely to be KFDV hosts in the crude bivariate analysis, however the relationship did not persist after accounting for feeding patterns and body mass. Neither were mass-corrected life history traits associated with infection status, suggesting a lack of allometric scaling of life history in this system. Other work has shown the fast-slow continuum of life history to be associated with diverging immune modulatory repertoires (Ricklefs and Wikelski, 2002). Along this continuum, fast-living species express higher reproductive output, reduced investment in reproductive output, and shorter lifespans (Molles, 2015). A corresponding decrease in adaptive immune function may make fast-living species more susceptible to infection, and thus increase the likelihood of their being a competent host (Johnson et al., 2012; Lee, 2006; Previtali et al., 2012). Alternatively, the likelihood of host competence may be enhanced among fast-living species by way of increased tolerance of infection burden (Råberg et al., 2009). In contrast to this specific ecological profile, the current study demonstrated a profile dominated primarily by size and feeding patterns alone. Furthermore, while life history traits did not scale with mass in the current study, other traits related to host competence also exhibit scaling and may be more relevant to host competence than life history in this system. For example, metabolism scales with body mass (Fossen et al., 2019; Kleiber, 1975), and can operate as a fundamental ecological constraint to many other biological parameters from individual species-level life history traits up to ecosystem-level processes (Brown et al., 2004). In addition, basal metabolic rate has been shown to influence host competence conjointly with body mass in tick-borne infections (Ostfeld et al., 2014). Unfortunately, as with species population density, sufficient data were not available to directly evaluate species basal metabolic rate on infection status in the current study.

This study has some limitations that warrant further comment. First, because of the long absence of animal field surveys investigating KFDV hosts, there are a limited number of infection positive species that could be included in this study. Moreover, the possibility of bias due to reporting effort cannot be ignored. We attempted to account for reporting effort first by controlling for the degree each species has been the subject of scientific study in the region, and second by restricting the analysis to only those species that have been the subject of virological study as a proxy for wildlife viral surveillance. While reporting effort exerted minimal influence on the results presented here, we should caution that the potential for residual bias cannot be entirely eliminated and therefore we emphasise the importance of renewed wildlife surveillance for KFDV across the region to validate the current findings. Second, some biological and life history characteristics exhibited substantial missing data for some species in the PanTHERIA database, which precluded their use in the analyses. Data imputation was employed where deemed appropriate (i.e. variables within a threshold of⍰<⍰70% missingness) using the methods described (Plourde et al., 2017; Stekhoven and Buhlmann, 2012), but the imputed data inevitably represent a diminished scope of mammalian life history in the Western Ghats.

## Conclusions

In conclusion we posit a preliminary ecological profile of KFDV wildlife hosts that emphasises consumption of plant resources as the main dietary strategy. Furthermore, KFDV hosts tended to be small, which could further contribute to transmission dynamics by influencing viral maintenance or amplification through the allometric scaling of as yet unidentified immune modulators. These ecological traits suggest the potential benefit of protecting primary forest habitat and its wildlife species as a means of interrupting transmission to humans. However, it is important to note that all the findings documented here are an initial attempt at an ecological profile that may help to guide new research efforts in KFDV wildlife surveillance, but will require validation from such efforts by way of comprehensive field investigations across the Western Ghats region.

## Acknowledgements

The authors received no funding for the conduct of this work.

