## Supplementary material for "A preliminary ecological profile of Kyasanur Forest disease virus hosts among the mammalian wildlife of the Western Ghats, India"

S1 Figure 1. Comparison of ecological and life history traits by taxonomic Order among the mammalian species of the Western Ghats.

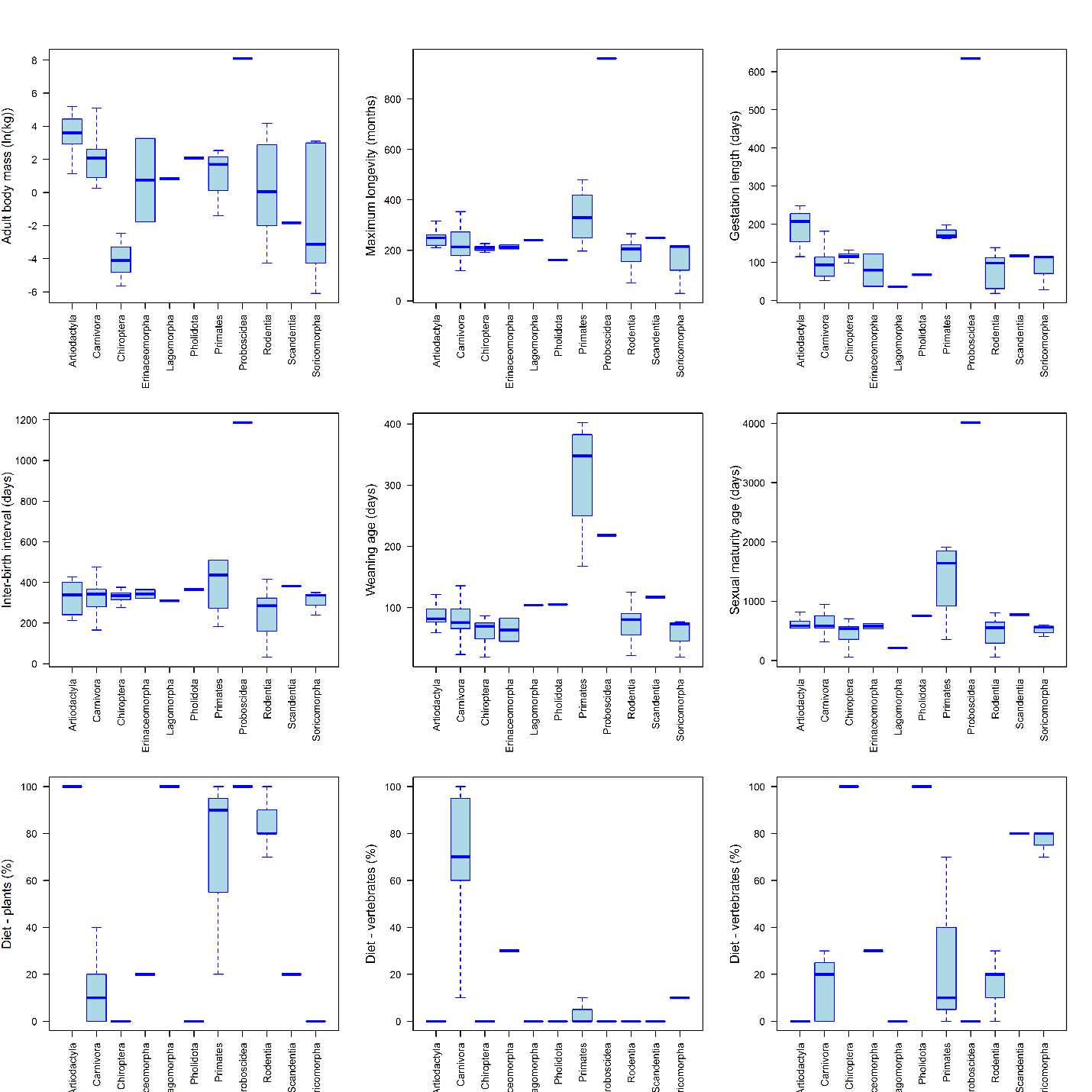

S2 Table 1. Blomberg’s K test for phylogenetic signal in species traits.

| Species trait | Blomberg’s K |
| --- | --- |
| Gestation length | 0.087 |
| Litter size | 0.079 |
| Maximum longevity | 0.057 |
| Weaning age | 0.069 |
| Inter-birth interval | 0.062 |
| Sexual maturity age | 0.077 |
| Body mass | 0.16 |
| Body mass gain | 0.14 |
| Diet – plants | 0.022 |
| Diet – invertebrates | 0.031 |
| Diet – vertebrates | 0.107 |
| Forage strategy | 0.087 |

S3 Table 2. Crude bivariate associations between species traits and host infection status derived from simple generalised additive models.

| Species trait | p-value |
| --- | --- |
| Body mass (ln(kg)) | 0.02 |
| Body mass gain | 0.64 |
| Maximum longevity (months) | 0.05 |
| Gestation length (days) | 0.86 |
| Inter-birth interval (days) | 0.03 |
| Litter size | 0.29 |
| Weaning age (days) | 0.13 |
| Sexual maturity age (days) | 0.05 |
| Diet – plants (%) | 0.0009 |
| Diet – vertebrates (%) | 0.12 |
| Diet – invertebrates (%) | 0.07 |
| Forage strategy | 0.05 |
| Circadian activity (Noctural) | 0.46 |
| Reporting effort | 0.06 |

S4 Table 3. **A**. Variance metrics for the factor loadings of the principal components analysis of the PanTHERIA life history traits: maximum longevity, sexual maturity age, gestation length, inter-birth interval, weaning age, and litter size. **B**. Correlation coefficients for each life history trait under the five factor loadings.

| A. Metric | PC1 | PC2 | PC3 | PC4 | PC5 |
| --- | --- | --- | --- | --- | --- |
| Standard deviation | 1.9552 | 0.9498 | 0.76765 | 0.58039 | 0.45096 |
| Proportion of variance | 0.6371 | 0.1504 | 0.11821 | 0.05614 | 0.03389 |
| Cumulative proportion of variance | 0.6371 | 0.7875 | 0.90571 | 0.94184 | 0.97574 |
| B. Life history trait | PC1 | PC2 | PC3 | PC4 | PC5 |
| Litter size | -0.246 | 0.889 | -0.251 | 0.111 | -0.242 |
| Sexual maturity age | 0.469 | 0.246 | -0.006 | 0.014 | 0.011 |
| Gestation length | 0.430 | -0.206 | -0.340 | 0.647 | -0.452 |
| Weaning age | 0.366 | 0.249 | 0.837 | 0.096 | -0.148 |
| Inter-birth interval | 0.440 | 0.011 | -0.251 | -0.744 | -0.369 |
| Maximum longevity | 0.455 | 0.211 | -0.240 | 0.077 | 0.761 |

S5 Table 4. Generalised additive model of host infection by mass-corrected life history (as represented by their principal component factor loadings) and dietary traits.

| Species trait | p-value* |
| --- | --- |
| Body mass-corrected life history (PC1) | 0.30 |
| Body mass-corrected life history (PC2) | 0.61 |
| Body mass-corrected life history (PC3) | 0.40 |
| Diet – plant (%) | 0.005 |

*p-values refer to the association between each species’ trait and infection status, not the overall fit of the model.

S6 Figure 2. The nonlinear relationships between Kyasanur Forest disease virus host probability and plant-based diet and body mass as derived from the best fitting generalised additive model restricted to only those wildlife species surveyed for viruses. Shaded areas represent the 95% confidence limits of the nonlinear function. Diet here represent the percentage of the species derived from plants and body bass is on the natural log scale.

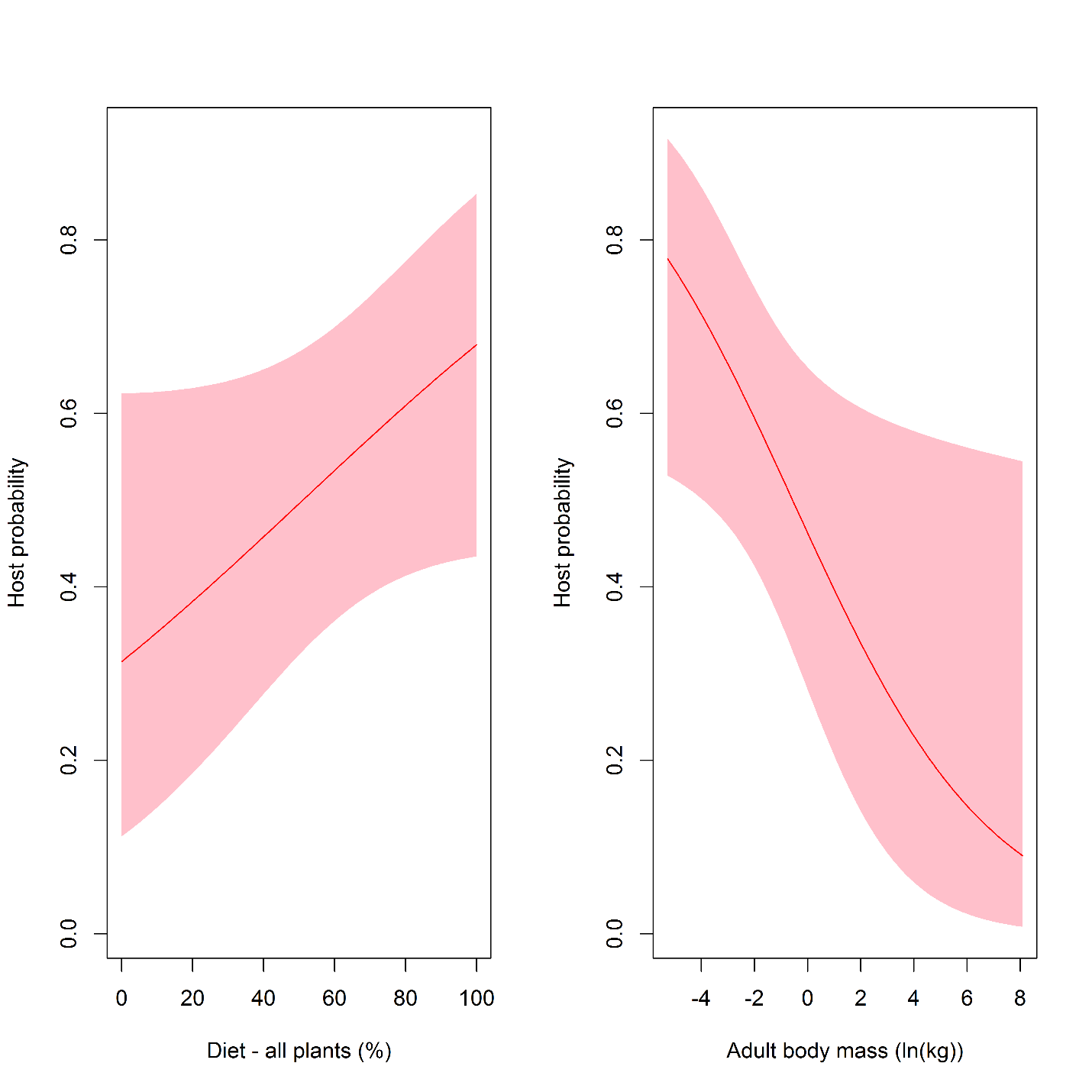
